# Rapid longitudinal SARS-CoV-2 intra-host emergence of novel haplotypes regardless of immune deficiencies

**DOI:** 10.1101/2021.12.22.473949

**Authors:** Laura Manuto, Marco Grazioli, Andrea Spitaleri, Paolo Fontana, Luca Bianco, Luigi Bertolotti, Martina Bado, Giorgia Mazzotti, Federico Bianca, Francesco Onelia, Giovanni Lorenzin, Fabio Simeoni, Dejan Lazarevic, Elisa Franchin, Claudia Del Vecchio, Ilaria Dorigatti, Giovanni Tonon, Daniela Cirillo, Enrico Lavezzo, Andrea Crisanti, Stefano Toppo

## Abstract

On February 2020, the municipality of Vo’, a small town near Padua (Italy), was quarantined due to the first coronavirus disease 19 (COVID-19)-related death detected in Italy. The entire population was swab tested in two sequential surveys. Here we report the analysis of the viral genomes, which revealed that the unique ancestor haplotype introduced in Vo’ belongs to lineage B and, more specifically, to the subtype found at the end of January 2020 in two Chinese tourists visiting Rome and other Italian cities, carrying mutations *G11083T* and *G26144T*. The sequences, obtained for 87 samples, allowed us to investigate viral evolution while being transmitted within and across households and the effectiveness of the non-pharmaceutical interventions implemented in Vo’. We report, for the first time, evidence that novel viral haplotypes can naturally arise intra-host within an interval as short as two weeks, in approximately 30% of the infected individuals, regardless of symptoms severity or immune system deficiencies. Moreover, both phylogenetic and minimum spanning network analyses converge on the hypothesis that the viral sequences evolved from a unique common ancestor haplotype, carried by an index case. The lockdown extinguished both viral spread and the emergence of new variants, confirming the efficiency of this containment strategy. The information gathered from household was used to reconstructs possible transmission events.

**AUTHOR SUMMARY:** It is of great interest and importance to understand SARS-CoV-2 ability to mutate generating new viral strains, and to assess the impact of containment strategies on viral transmission. In this study we highlight the rapid intra-host haplotype evolution regardless of symptom severity and immune deficiencies that we observed during the first wave of the pandemic in the municipality of Vo’ in Italy. The confirmation that all the haplotypes found in this small community derive from a common ancestor haplotype, has allowed us to track the rapid emergence of new variants but lockdown and mass testing efficiently prevented their spread elsewhere.

## INTRODUCTION

Severe acute respiratory syndrome coronavirus 2 (SARS-CoV-2) infection continues to spread world-wide, with over 5 million of deaths out of more than 250 million of positive cases reported since the beginning of the pandemic (1). There is extensive evidence that SARS-CoV-2 genome has evolved and has acquired several mutations conferring higher viral fitness, with almost all genomic sites being affected by mutation events (2–4).

Here we provide novel insights into the generation of diversity of SARS-CoV-2 genomes in a close community.

On February and March 2020, preceding and following a two-weeks lockdown respectively, two mass swab testing campaigns were conducted in Vo’ to trace and isolate all the positive subjects. In previous work, we investigated the effectiveness of the implemented non-pharmaceutical interventions (NPIs), the role of asymptomatic infections (5) and the tracking of T-cell signatures of the whole population (6). In May and November 2020, two follow-ups were carried out to assess the antibody dynamics following infections (7).

Here we report on the results from the sequencing of SARS-CoV-2 genomes extracted from oro-nasopharyngeal swabs of positive subjects. We obtained the sequences for 87 samples, representing the vast majority of the positive subjects detected in Vo’. Sequencing data has allowed us to identify unequivocally the ancestor haplotype of the virus circulating in Vo’ at the time of the outbreak. This evidence provided the opportunity to infer the evolution of SARS-CoV-2 in an isolated community, from an intra-host, a household and a general perspective, tracing the generation of novel mutations from the index haplotype. Information on date of symptoms onset allowed us to reconstruct a temporal expanding network of infected subjects. The uniqueness of this study lies in the feature of this isolated and small community that, since the very beginning of the SARS-CoV-2 pandemic, was monitored over time, enabling us to investigate the virus evolution and transmission dynamics.

## RESULTS

### Viral haplotypes circulating in Vo’

During the first surveys, we collected oropharyngeal swab samples of nearly the entire population of Vo’ at two consecutive time points, (21-29 February 2020 and 7 March 2020, respectively). Out of the tested subjects, 100 individuals had a positive result, with 12 of them testing positive again at the second time point. We sequenced the viral genome of all the available samples. As a result, 87 consensus sequences were obtained and uploaded in GISAID (8).

All the viral sequences were characterised by two mutations, *G11083T* and *G26144T* (from here on reported as Ancestor Haplotype, AH). Interestingly, these mutations were previously identified in a couple of Chinese tourists who, after visiting Verona, Parma and Florence, were diagnosed with SARS-CoV-2 infection in Rome on 31^st^ January 2020, and correspond to the first reported case of COVID-19 in Italy (9). Surprisingly, this haplotype, which is classified as lineage B according to Rambaut’s nomenclature (10), accounted for 43.8% of total sequences in the February-March 2020 in the Veneto region. In the surrounding regions, such as Lombardia and Trentino Alto Adige (11), we found evidence of a different source of introduction, given that the B.1 lineage was dominant (12) (Fig 1).

**Fig 1.**
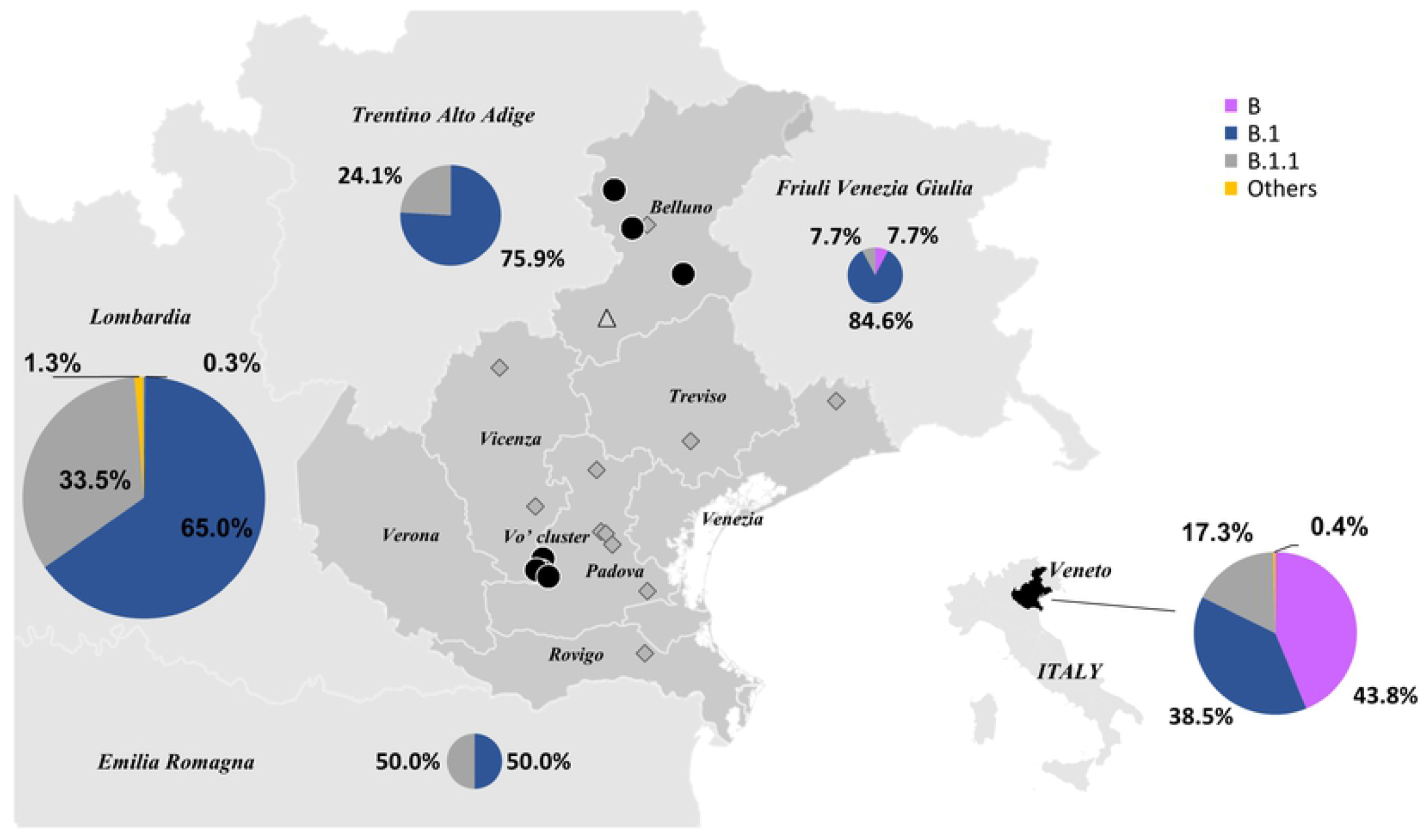
Lineages circulating in Northern Italian regions. Percentages of SARS-CoV-2 lineages circulating in Emilia Romagna (n=2), Friuli Venezia Giulia (n=13), Lombardia (n=400), Trentino Alto Adige (n=58) and Veneto (n=226) regions in February and March 2020 according to GISAID database. The size of the pie charts reflects the number of sequences available per region. The different lineages are colored according to the legend. Black circles represent haplotypes identical to those found in Vo’, diamonds indicate haplotypes with one additional mutation (edit distance +1 from Vo’) and triangles correspond to haplotypes with two additional mutations (edit distance +2 from Vo’).

While the AH was found only in Vo’ and in the province of Belluno, some AH’s derivative subtypes were observed in other provinces of Veneto region.

The sequencing data indicate that the B lineage haplotypes introduced in the Veneto region, as the ones identified in Vo’, derive from the AH, carrying G11083T and G26144T mutations, that further developed new additional mutations.

According to the sequenced data, the AH was the most prevalent in Vo’ (38 out of 87, 43.7%), while additional 26 unique haplotypes derive from the ancestor one. A total of 42 unique point mutations defining the different haplotypes, (including *G11083T* and *G26144T*) mapped along different regions of the viral sequence, 1 was located in 5’-UTR (2.4%), 15 were synonymous (35.7%) and 26 were non-synonymous (61.9%). Moreover, a 9-nucleotide deletion was observed in two samples, resulting in 3 amino acid deletion affecting the non-structural protein 1 (nsp1) (*ORF1a*:*K141*-,*ORF1a*:*S142*-,*ORF1a:F143-*), a mutation already observed in other sequenced SARS-CoV-2 genomes belonging to different lineages (13) and different countries, suggesting homoplasy events.

Although the short time transmission window hampered the detection of a natural or purifying selection, most of the non-synonymous mutations found in Vo’ emerged independently later on during the pandemic in different countries (S1 Table). The two mutations defining the AH behaved differently during the pandemic. *G26144T* was mostly associated with *G11083T* in the lineage B and disappeared in April 2021 after reaching a peak at the beginning of the pandemic (February-April 2020). Conversely, the *G11083T* mutation appears in recently collected sequences belonging to different lineages, suggesting to be another example of homoplasy (14) (S1 Fig).

Interestingly, the mutation *Spike:L5F* (*C21575T*), characterising the viral haplotype of two subjects in Vo’, emerged from January 2021 to August 2021 as part of the mutation pattern defining the B.1.526 lineage (Iota variant), prevalent in USA (15) (S2 Fig).

### Vo’ haplotypes in Europe

We compared the viral genomes collected at Vo’ with a selection of contemporary European sequences from different locations. At the beginning of the pandemic the AH was circulating also in other European countries, with UK, Italy (90% in Veneto region) and Spain reporting the highest occurrences. To analyse the data, we used two different approaches based on phylogenetic analysis (Maximum Likelihood phylogenetic tree, ML) and on the construction of a Minimum Spanning Network, MSN (see Methods).

In Fig 2, we provide a ML phylogenetic tree and a MSN limited to Vo’ haplotypes and to closely related European haplotypes, where all the European sequences carrying the AH, including the Italian ones, are represented as a big green node in the MSN, while they are collapsed in the phylogenetic tree. The same analysis, extended to all the European sequences collected at the time period considered, are provided in S3 and S4 Figs for the phylogenetic tree and the MSN, respectively. According to phylogenetic and MSN reconstruction, only one sequence, “*VO_SR_65*” (*haplotype G11083T*, *G26144T*, *C22088T*), shares a private mutation with a Polish sequence, *EPI_ISL_455452* (*haplotype C4338T*, *T9743C*, *G11083T*, *G26144T*, *C22088T*). However, the Polish sequence (dated 2020-03-28) relates to a sample collected one month later than the Vo’ sample “*VO_SR_65*” (dated 2020-2-21), suggesting an independent evolution and possibly a homoplasy event.

**Fig 2.**
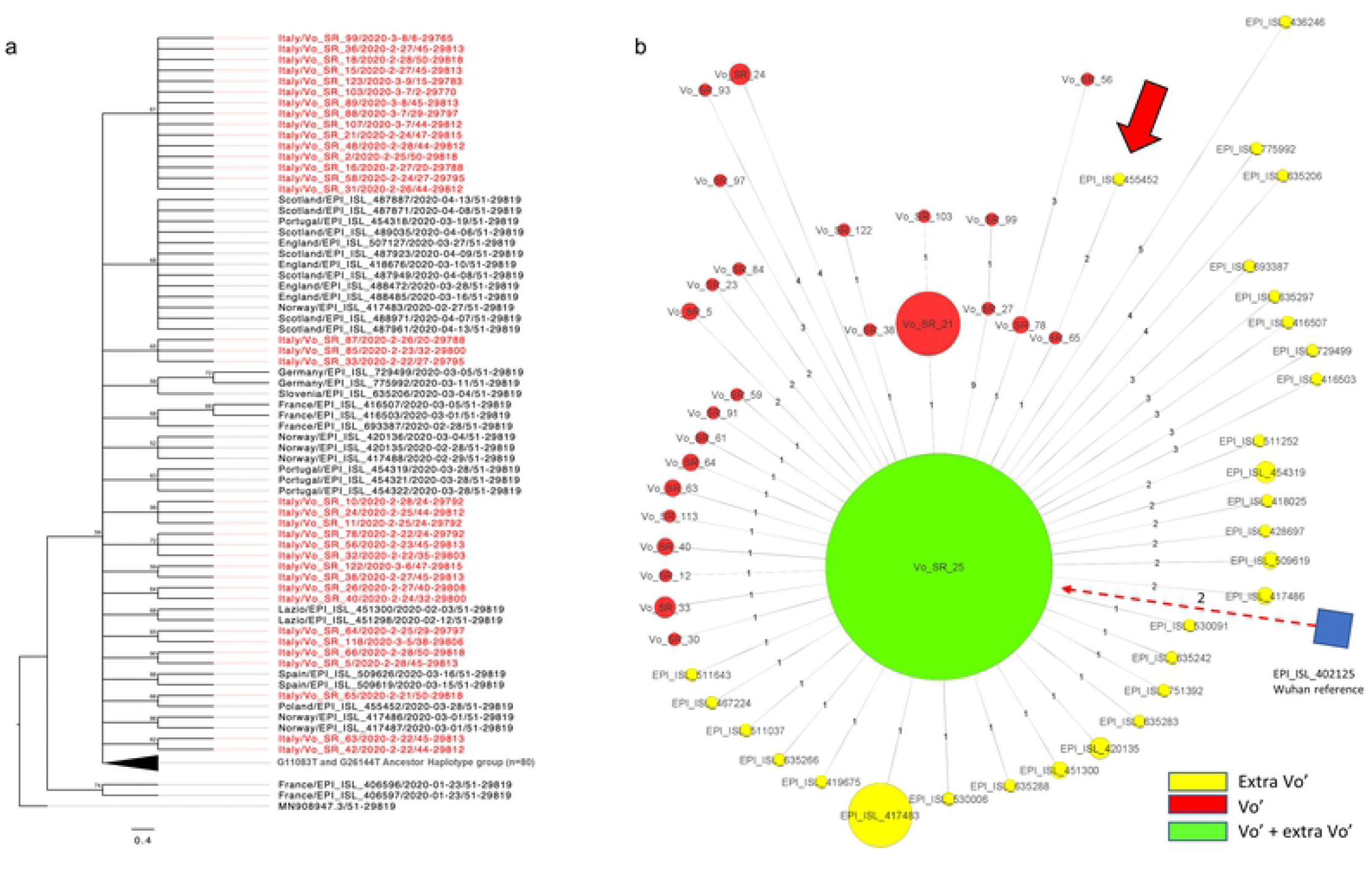
Maximum Likelihood phylogenetic tree and Minimum Spanning Network of sequences related to Vo’ viral genomes. European sequences closely related to Vo’ viral genomes and with collection date corresponding to the beginning of the pandemic were retrieved from GISAID and utilised for the ML phylogenetic tree and MSN.

a. Maximum Likelihood phylogenetic tree. The phylogenetic tree is collapsed in correspondence of the AH, found both in Vo’ and other European countries. Sequences from Vo’ are reported in red.
b. Minimum Spanning Network. The AH is represented as a big green node, containing both Vo’ and European sequences. The size of the nodes reflects the abundance of identical sequences. The distance of the edges from the central node reflects the number of accumulated mutations. European/Italian sequences are reported in yellow, Vo’ sequences are reported in red, and Wuhan reference node is represented as a blue square. The only extra-Vo’ haplotype genetically related to a Vo’ haplotype is indicated by the red arrow.

These results confirmed that the Vo’ haplotypes were not observed in other countries and did not generate descendants, as a consequence of mass testing and lockdown strategy implemented in Vo’(16–18).

### Intra-host viral evolution

We obtained the viral sequence of 12 individuals positive at two sequential swab tests. As reported in Fig 3, in 4 out of 12 subjects (33.3%) the virus acquired at least a novel mutation in an average time of 11 days (range 9-16 days). While 3 out of 4 individuals acquired a single mutation at the second time point, one subject (samples *VO_SR_7* and *VO_SR_93*) accumulated 4 mutations in 13 days, highlighting the rapidity of viral evolution in some subjects. No significant differences in days after the first swab test, disease severity, antibody production or T cell receptor breadth or depth (6) were found comparing individuals with a constant viral haplotype to individuals with a mutated viral haplotype (Figs S5 and S6). Giving that all the haplotypes identified in Vo’ derive from the AH, and that within approximately two weeks of positivity, the viral genomes accumulated at least a novel mutation in 33.3% of cases, we further investigated this event among all the available subjects. Considering that in Vo’ we identified 26 unique haplotypes (AH excluded) out of 87 subjects and the two-weeks lockdown imposed by the local authorities, we can assume that intra-host evolutionary events occurred in 26 subjects out of 87 (29.9%), within this time window. Of these 26 cases, 4 were proven by the sequential swab tests, as described above, while other 10 cases could be inferred according to viral haplotype evolution within family households, virus haplotypes and the symptoms onset dates (Table 1).

**Fig 3.**
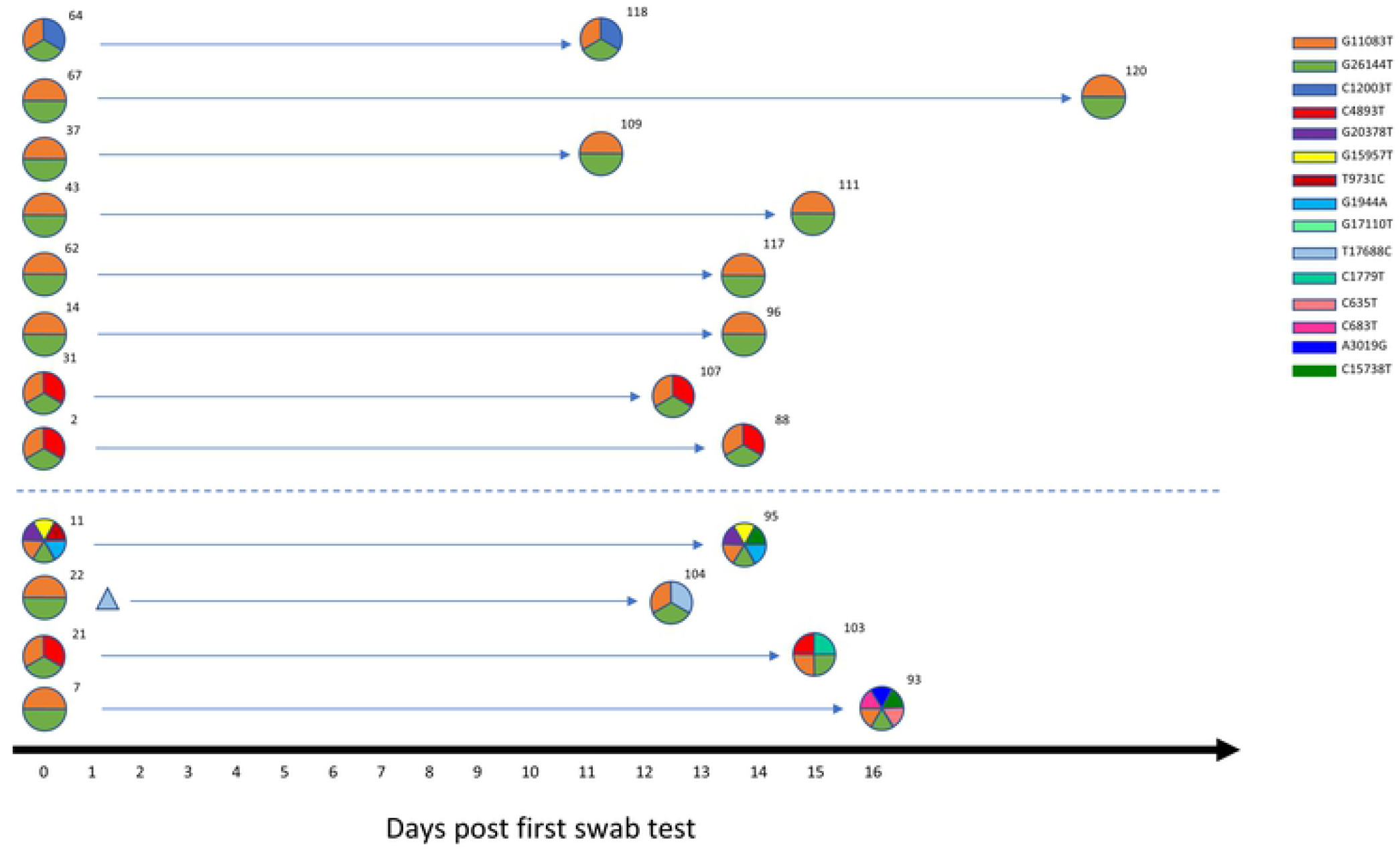
Within host variation. Longitudinal analysis of SARS-CoV-2 intra-host variation in 12 subjects in an average of 11-day time window. The viral haplotypes of the two consecutive swab tests of each subject are reported on the same line. Each haplotype is depicted as a circle with each slice representing a mutation, characterised by a colour according to the legend. Minor variants are represented as triangles.

**Table 1.**
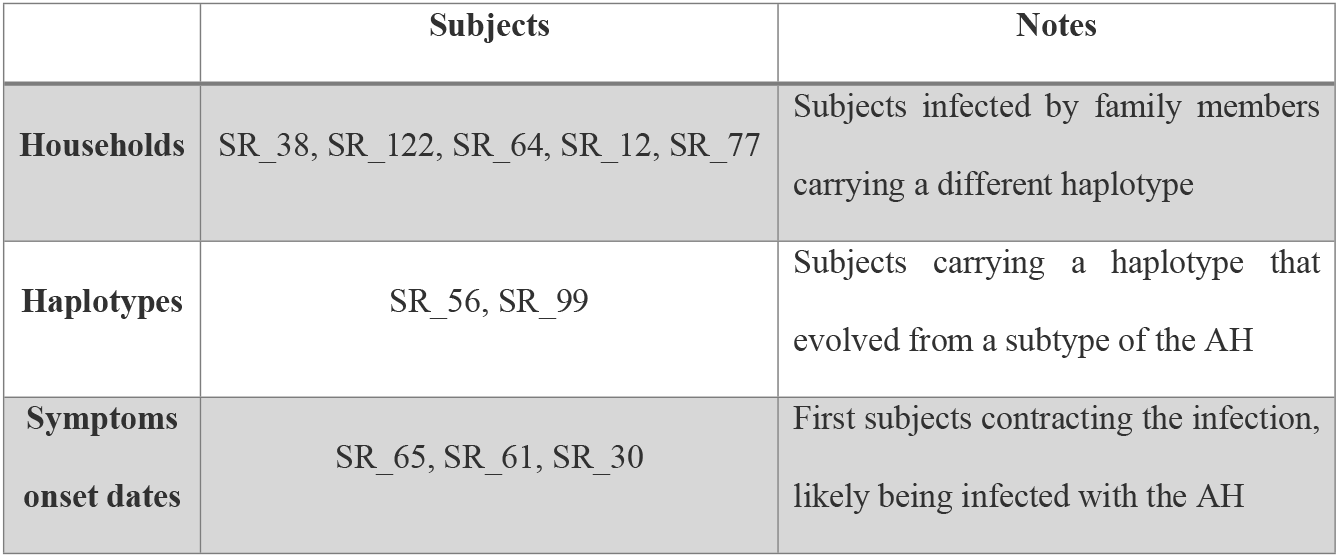
Inferred intra-host viral evolution.

|  | Subjects | Notes |
| --- | --- | --- |
| **Households** | SR_38, SR_122, SR_64, SR_12, SR_77 | Subjects infected by family members carrying a different haplotype |
| **Haplotypes** | SR_56, SR_99 | Subjects carrying a haplotype that evolved from a subtype of the AH |
| **Symptoms onset dates** | SR_65, SR_61, SR_30 | First subjects contracting the infection, likely being infected with the AH |

### Within-household variability

The different haplotypes identified in Vo’ were analysed in the context of the household structure to investigate potential transmission chains and within household viral evolution.

The intra-family transmission chains reported in Fig 4 were based on: a) swab test results and relative dates, b) viral consensus genome, c) minor variants, considered as alternative nucleotides with a frequency from 5% to 49%. We assumed that the subjects who negativized earlier were the infectors and assumed no reversion.

**Fig 4.**
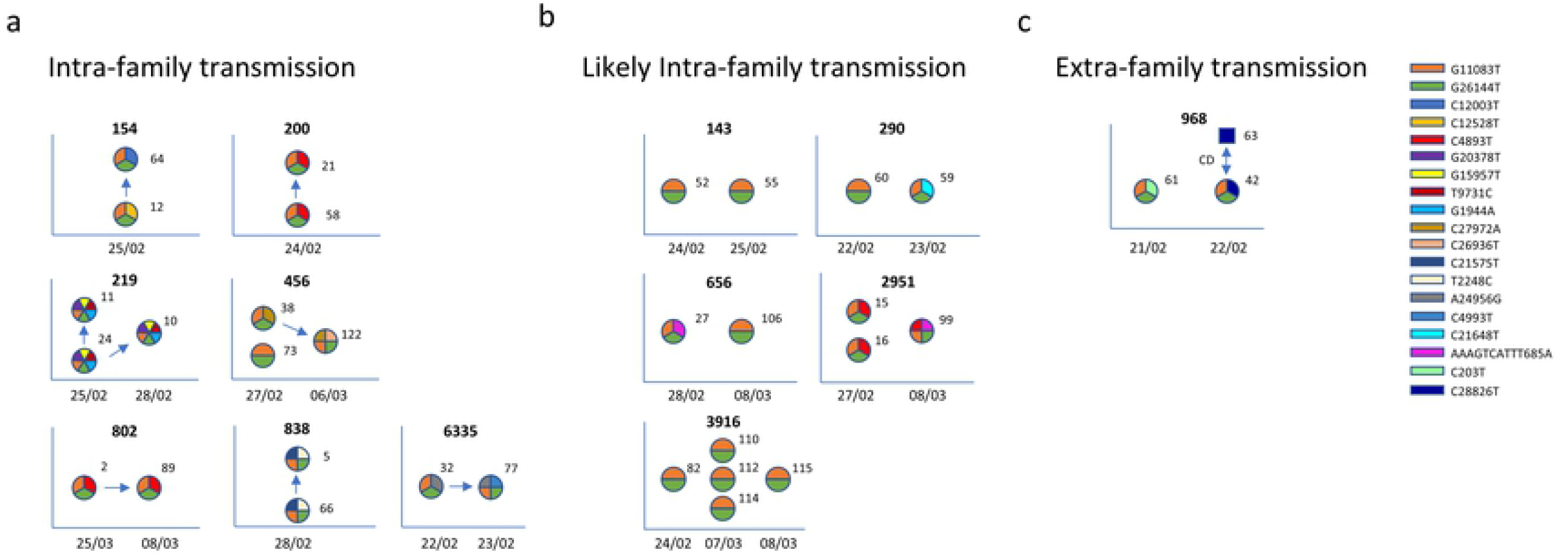
Within family diversity and transmission. Sequences were grouped according to the households and analysed separately. Each haplotype is depicted as a circle with each slice representing a mutation, characterised by a colour according to the legend. For each household (identification number provided at the top of each diagram) genetic information of family members (labelled with an identification number) and relative metadata (date of positive swab tests reported on x axis) were utilised to investigate the diversity and the transmission chain. Households were then subdivided into within household transmission (panel a), uncertain transmission (panel b), and extra-household transmission (panel c).

Interestingly, we found evidence that a few mutations were household specific (not present in any other sample collected in Vo’ outside the housed). Households 838 and 219, where the 2 out of 2 and 3 out of 3 family members shared the same haplotype (*G11083T*, *G26144T*, *T2248C*, *C21575T*, and *G11083T*, *G26144T*, *G1944A*, *T9731C*, *G15957T*, *G20378T* respectively), exemplify this phenomenon.

Household 456, where one household member is characterised by the AH (haplotype *G11083T*, *G26144T*), the second one acquired a novel private mutation (haplotype *G11083T*, *G26144T*, *C27972A*) and the third one further acquired another private mutation (haplotype *G11083T*, *G26144T*, *C27972A*, *C26936T*), likely reflects the direction of transmission and the sequential viral evolution during transmission among household members.

By enriching genetic data with metadata, such as swab tests and contacts information, it was possible, for 8 out of 13 (61.5%) households, to reconstruct the household transmission chains, depicted with arrows in Fig 4 panels a and b. Notably, while 7 out of 8 of the reconstructed transmissions occurred within the same household Fig 4a, in one case we found evidence of household members contracting the infection outside of the household. This is confirmed by the difference in viral haplotypes between the two household members, with one of them sharing the same haplotype with a declared direct contact Fig 4c. In the remaining 5 households, although the transmission probably occurred within family members, we did not have enough information to reconstruct the transmission chains (Fig 4b).

### Transmission chain reconstruction

In Fig 5, contact (panel on the left) and genetic (panel on the right) data are reported and compared. Given the incompleteness of the contact information and of the reported dates of symptom onset, it was not possible to identify unequivocally the chains of transmission events in Vo’, not even with the support of genetics. However, the data suggest that the outbreak initiated from an indoor space that fuelled a superspreading event. The time lapse animation (see S7 movie) shows the reported contacts according to the reported times of symptoms onset.

**Fig 5.**
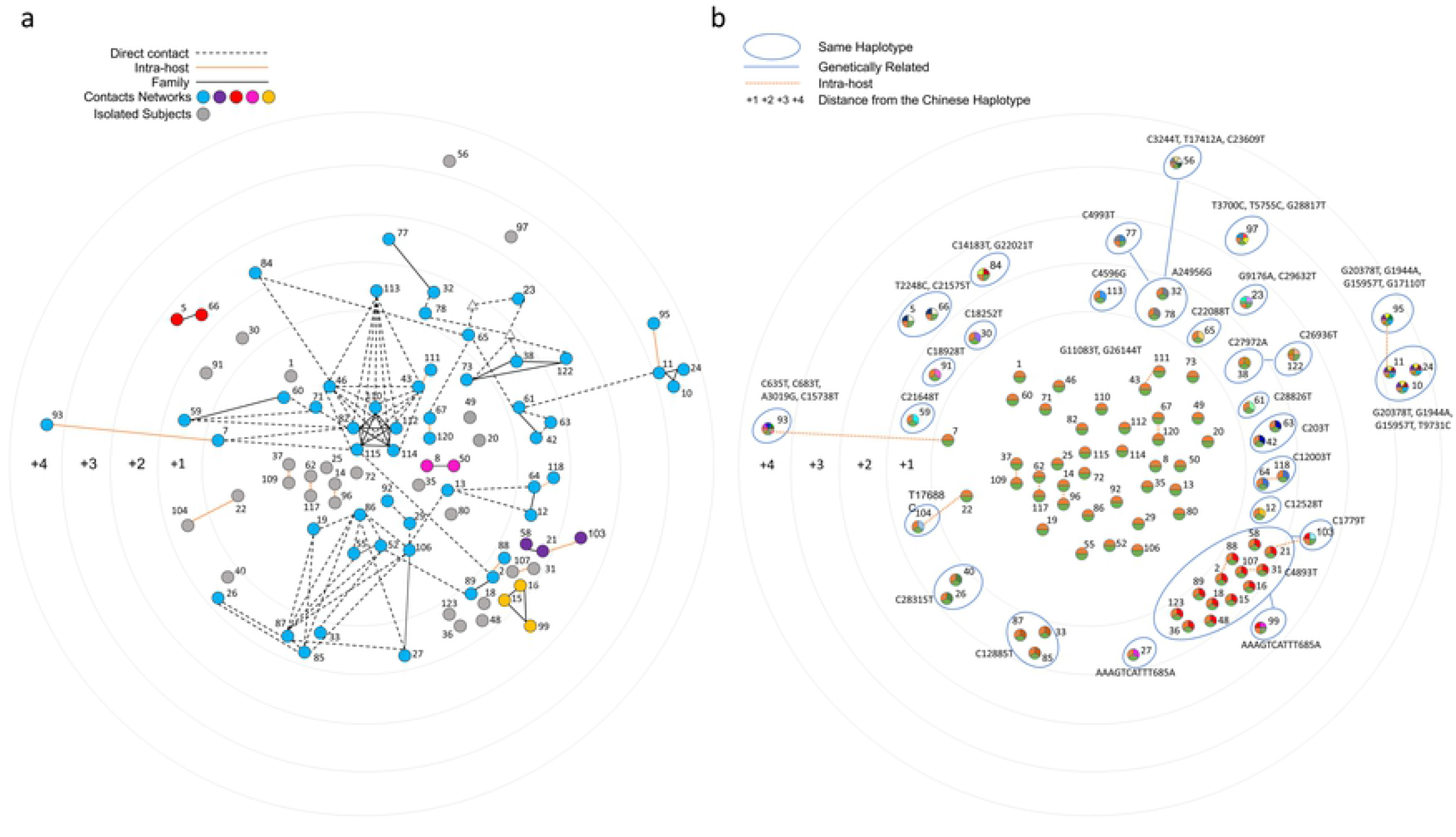
Network of contacts based on post-infection interviews. a) samples are connected according to the contacts declared during the post-infection interviews. Each contacts chain is coloured according to the legend, with nodes not declaring any contact being coloured in grey. Subjects who declared informative contacts, but without an available viral sequence, are depicted as white triangles. b) Vo’ haplotypes are clustered according to their mutations and to the genetic distance from the AH (edges at distance 1 are not drawn for graphical reasons)

## DISCUSSION

The population of Vo’ was the first in Italy and Europe to be placed under lockdown by the regional authorities after the first ascertained COVID-19 death in the country. This situation has allowed us to study in details different aspects of the epidemic in a controlled epidemiological setting.

The sequencing data identified the Vo’ Ancestor Haplotype (AH), characterised by *G11083T* and *G26144T* mutations, from which all the other sequenced haplotypes evolved. This haplotype, initially observed in two Chinese tourists visiting Italy in January 2020, was dominant in Veneto region at the beginning of the pandemic, while lineage B.1 was prevalent in the surrounding regions, revealing multiple viral introductions in Italy (19).

Moreover, MSN and phylogenetic analysis including European sequences of the time confirmed that, although the AH was circulating also in other countries, none of the descendant haplotypes evolved in Vo’ was observed elsewhere. Thus, the lockdown was efficient in containing the outbreak in Vo’ and the new virus variants emerged from it. Interestingly, we observed three cases of putative homoplasy, with: a) *C22088T*, present in a Polish sequence collected one month later than Vo’ outbreak, b) *G11083T*, found in several sequences from different countries and lineages, c) *C21575T*, shared by two Vo’s household members and appearing at a global level only in 2021, as a mutation defining the B.1.526 lineage.

According to present literature, the accumulation of viral mutations at the consensus level was previously observed in treated (20,21) or immunocompromised subjects (22,23) experiencing prolonged infections whilst other studies reported only longitudinal changes of intra-host minor variants (24,25). We here provide the first evidence of the rapid intra-host accumulation of mutations at the consensus level in a time window as short as two weeks regardless of symptoms severity or immunodeficiencies. The generation of new haplotypes occurred in average time of 11 days in 33% of samples (4 out of 12) in otherwise healthy individuals.

In an exemplary case, a total of four mutations accumulated in 13 days, pointing out the rapidity of viral evolution. Based on the collected data, we could infer a frequency of intra-host evolution of 30% (26 out of 87). Although no correlation with symptoms severity, length of the infection and serological data was found, other mechanisms could be responsible for within-host viral evolution, such as host immune response (26,27).

Given the frequency of intra-host viral evolution, it was difficult to define a transmission chain based on genetics, since the differences in the haplotypes can derive either from the haplotype carried by the infector or by an intra-host evolution occurred in the infectee. The integration of sequencing data with metadata allowed us to infer the transmission chains among some household members indicating within household transmissions as the most frequent ones (7 out of 13).

Though contact tracing analysis was limited to symptoms onset data to reconstruct a temporal expanding network of infected subjects, it enabled us to identify the first infections and a gathering place which had a main role in the transmission of the infection.

To sum up, this study sheds light around the effectiveness of contact tracing based on interviews, which showed limitations in the coverage and traceability of the contacts among the identified infections. For these reasons, in the absence of effective digital contact tracing, mass testing followed by case isolation represents the best approach, when applicable, for identifying and isolating infections thus giving the best chances to achieve epidemic control (28–30). As a matter of fact, the variants generated in the Vo’ population, did not spread outside Vo’ showing that effective control strategies not only curb transmission, but also control the emergence and spread of new variants. The number of mutations observed in the Vo’ population over just two weeks should warn about the velocity of adaptation that may be occurring in different subjects and the potential ability of SARS-CoV-2 to develop immune evasion.

## MATERIALS AND METHODS

### Sample collection, library construction and sequencing methods

The presence of SARS-CoV-2 genomes was evaluated with an in-house real-time RT–PCR method targeting the envelope gene (E) (31). Total nucleic acids were purified from nasopharyngeal swab samples using a MagNA Pure 96 System (Roche Applied Sciences). All samples were treated with DNase I (Promega, Madison, WI, USA) to eliminate residues, of the host, and bacterial gDNA. Quantity and quality checks were performed on the HSRNA chip for Tapestation 4100 (Agilent, Santa Clara, CA, USA). Samples with low RNA concentration were concentrated using Vacufuge plus (Eppendorf, Hamburg, Germany). Starting with 50 ng of total RNA, NEBNext Ultra II First Strand and Non-Directional RNA Second Strand Synthesis Module (NEB, Ipswich, Massachusetts) were used to convert RNA into cDNA. Before proceeding with the next steps, we performed RTqPCR on all samples in order to determine the relative abundance of viral RNA. RTqPCR was performed by amplification regions for the following target genes: RNA-dependent RNA-polymerase (RdRP), nucleocapsid (N), envelope (E), and Internal Control.

The enrichment step was performed according to the manufacturer instructions. We used the Illumina Nextera Flex kit for Enrichment/Respiratory Virus Oligos Panel Detection of SARS-CoV-2 RNA. Briefly, samples were fragmented and barcoded with Nextera Flex, followed by PCR amplification (17 cycles). Before hybridizations with Respiratory Virus Oligos Panel, samples were pooled according to the viral load (previously determined by RTqPCR). Each pool was composed by 8 samples with similar Ct (threshold cycle), to avoid bias competition between samples with low and high Ct during hybridization and sequencing.

After a quality check on HSDNA chip for Tapestation 4100(Agilent), libraries were diluted to 2.2pM, mixed with Pfix (20%) and then loaded on NextSeq 500 sequencher (Illumina, San Diego, CA). Sequencing was performed in PE (pair-end) mode, generating 149nt length reads, and 1-5M clusters for each sample.

### Viral genome assembly: quality check and mapping of the reads

Raw sequences were filtered for length and quality with Trimmomatic v0.40 according to the following parameters: ILLUMINACLIP:TruSeq3-PE-2:2:30:10 LEADING:30 TRAILING:30 SLIDINGWINDOW:4:20 MINLEN:90. High quality reads were aligned on the SARS-CoV-2 reference genome (genbank ACC: NC_045512) with BWA-MEM v0.7.1. Duplicated reads were then removed with Picard tool v2.25.0 (http://broadinstitute.github.io/picard/). Consensus sequence were generated using a combination of SAMtools v1.11 and VarScan v2.4.1 variant caller. Consensus sequences were reconstructed from VarScan output with an in-house script that automatically introduces ‘N’ in low quality or uncertain/uncovered regions of the reference sequence.

### Sequence selection from GISAID

The Vo’ genome sequences of SARS-CoV-2, divided by lineage, were used as input of cd-hit-est-2d (32) and compared with the GISAID database. The first 20 most similar genomes of every cluster were used as input of MAFFT (33) to produce a multiple alignment with the Wuhan genome (GenBank ID MN908947.3) as reference. This dataset has been used to build the minimum spanning network (MSN) and the phylogenetic tree.

### Minimum spanning network (MSN)

The MSN(34) was built to find out the most probable path of contagion starting from a distance matrix calculated by the genome sequences of lineage B as classified by the Pangolin tool (https://github.com/cov-lineages/pangolin). The nodes of the MSN represent the genomes that share the same pattern of SNPs (with reference to the Wuhan genome MN908947.3), while an edge connects two nodes if they have compatible genomes (e.g. one can be obtained by the other by adding some mutations). Nodes are labelled with the sampling location and date of the corresponding sequences, edges with the number of mutations that differentiate the two sets of connected sequences. The minimum spanning network algorithm used to reconstruct the MSN from the distance matrix connects the virus sequences minimizing the edit distance (Hamming distance). In the reconstructed network, we defined a possible introduction as a node containing also sequences sampled outside the Vo’ area. Additionally, the sampling date of the Vo’ node (i.e. the introduction) must be later than that of the other GISAID sequences. Similarly, a possible exit is a node containing only Vo’ sequences that are the source of an edge leading to a node containing only GISAID sequences (i.e. and no sequence from Vo’). Moreover, the sampling date of the Vo’ node must precede that of the GISAID sequences. The MSNs were finally pruned to remove nodes not directly connected to a node containing sequences from Vo’.

### Phylogenetic analysis

Two different sequence sets were used for depicting the phylogenetic relationships between Vo’ samples and the European epidemiological framework. The first samples set included a total of 1252 full genome sequences from Europe, collected in the same pandemic period. The second data set included a selection of sequences from the first set (n = 148). Both sets included SARS-CoV-2 Wuhan-Hu-1 isolate (Genbank accession number MN90894) as reference (GenBank accession number: MN908947.3). The 90 bp end at both 5’ and 3’ were eliminated, to avoid possible sequencing errors. *Phylogenetic* trees were constructed using RAxML (35) software. Each tree is statistically supported by bootstrap process (100 replicates) and Wuhan isolate was used as outgroup.

## ETHICAL APPROVAL STATEMENT

The study was approved by the Ethics Committee for Clinical Research of the province of Padua under protocol number 0022591 - 7/4/2020. Study participation was by consent. For participants under 18 years of age, consent was provided by a parent or legal guardian.

## ACKNOWLEDGEMENTS

This work was supported by the Veneto Region. E.L. and S.T. acknowledge research funding from the European Union’s Horizon 2020 research and innovation programme, under grant agreement No 874735 (VEO). E.L. acknowledges funding from the University of Padova and the Department of Molecular Medicine (STARS-CoG ISS-MYTH and PRID/SID 2020). S.T. acknowledges funding from the University of Padua (TOPP_PRIV20_01 and TOPP_SID19_01). G.T. and A.C. acknowledge funding from Fondazione Umberto Veronesi, Misura Ricerca Covid 19, year 2020. We gratefully acknowledge the Authors from the Originating Laboratories and the Submitting Laboratories who generated and shared via GISAID the data on which part of this research is based.

## SUPPORTING INFORMATION

**S1 Fig.**
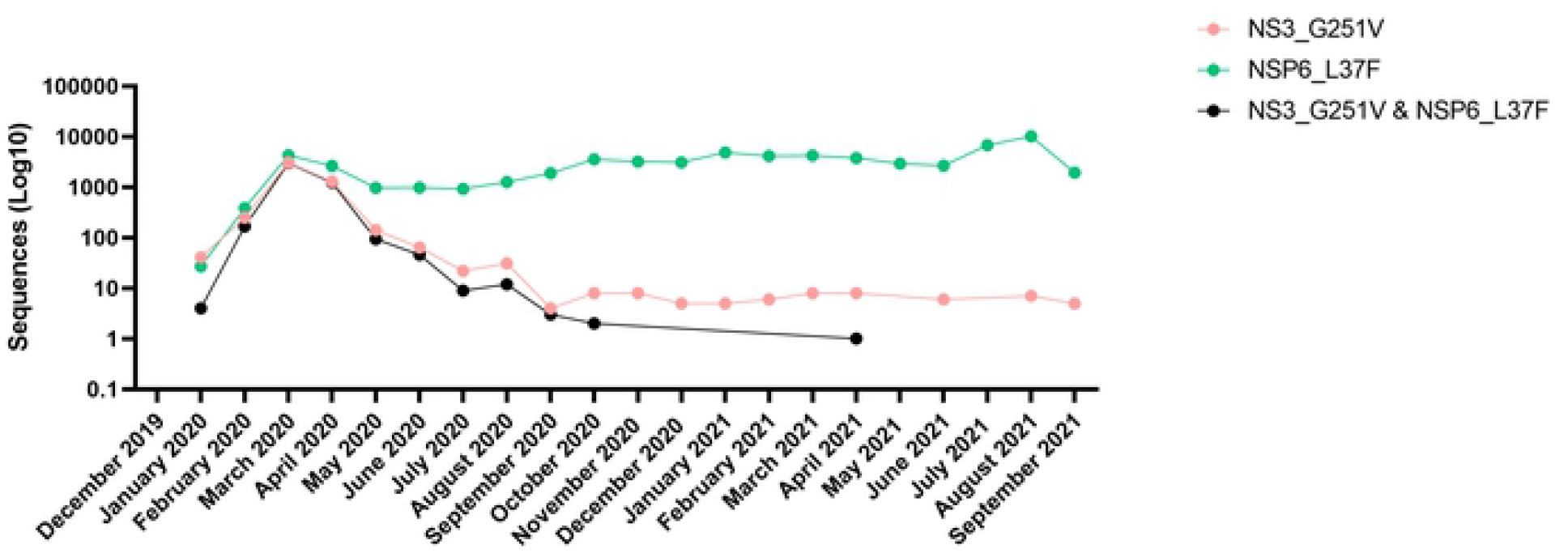
Mutations defining Vo’ haplotype in time. Number of sequences carrying G11083T (NSP6:L37F) and G26144T (NS3:G251V) mutations deposited in the GISAID database from December 2019 to September 2021. Mutations were analysed separately (green and pink line) and as part of the same haplotype (black line).

**S2 Fig.**
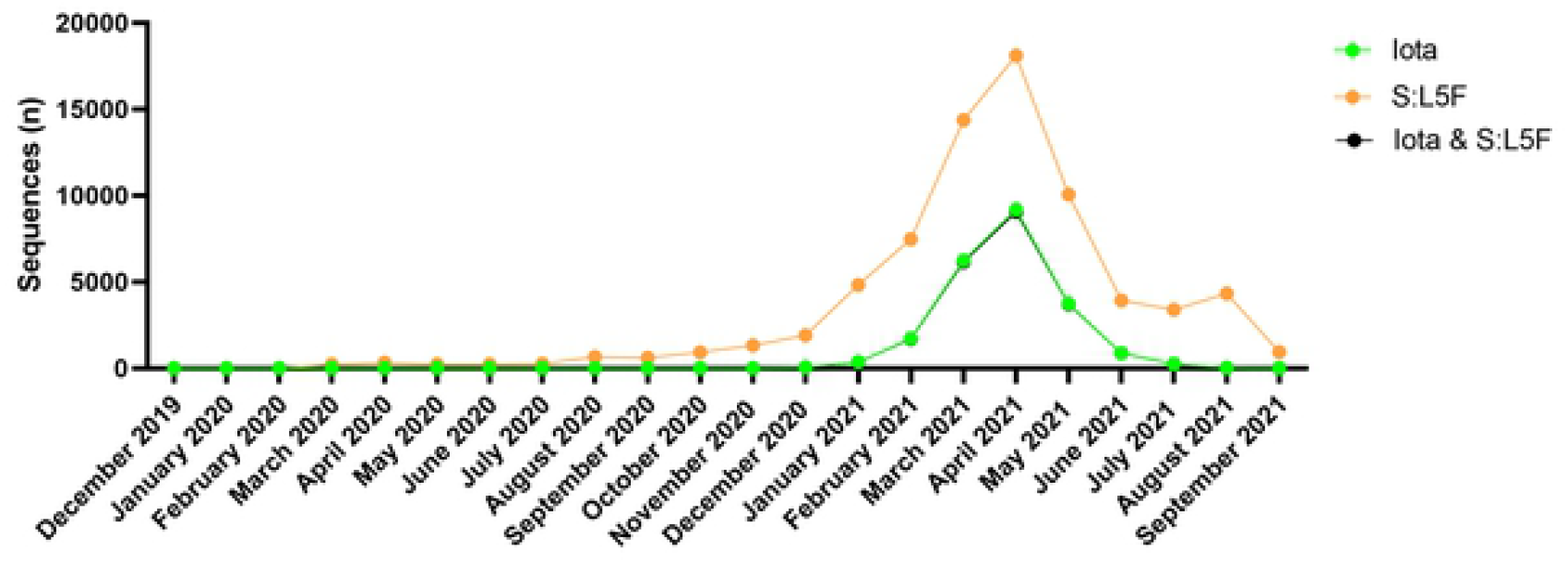
S:L5F appeared in Iota variant in 2021. Number of sequences deposited in the GISAID database and collected from December 2019 to September 2021 carrying C21575T (S:L5F) mutation (yellow line) compared to sequences belonging to 21F clade (Iota variant, green line) and the combination of the two cases (black line).

**S3 Fig.**
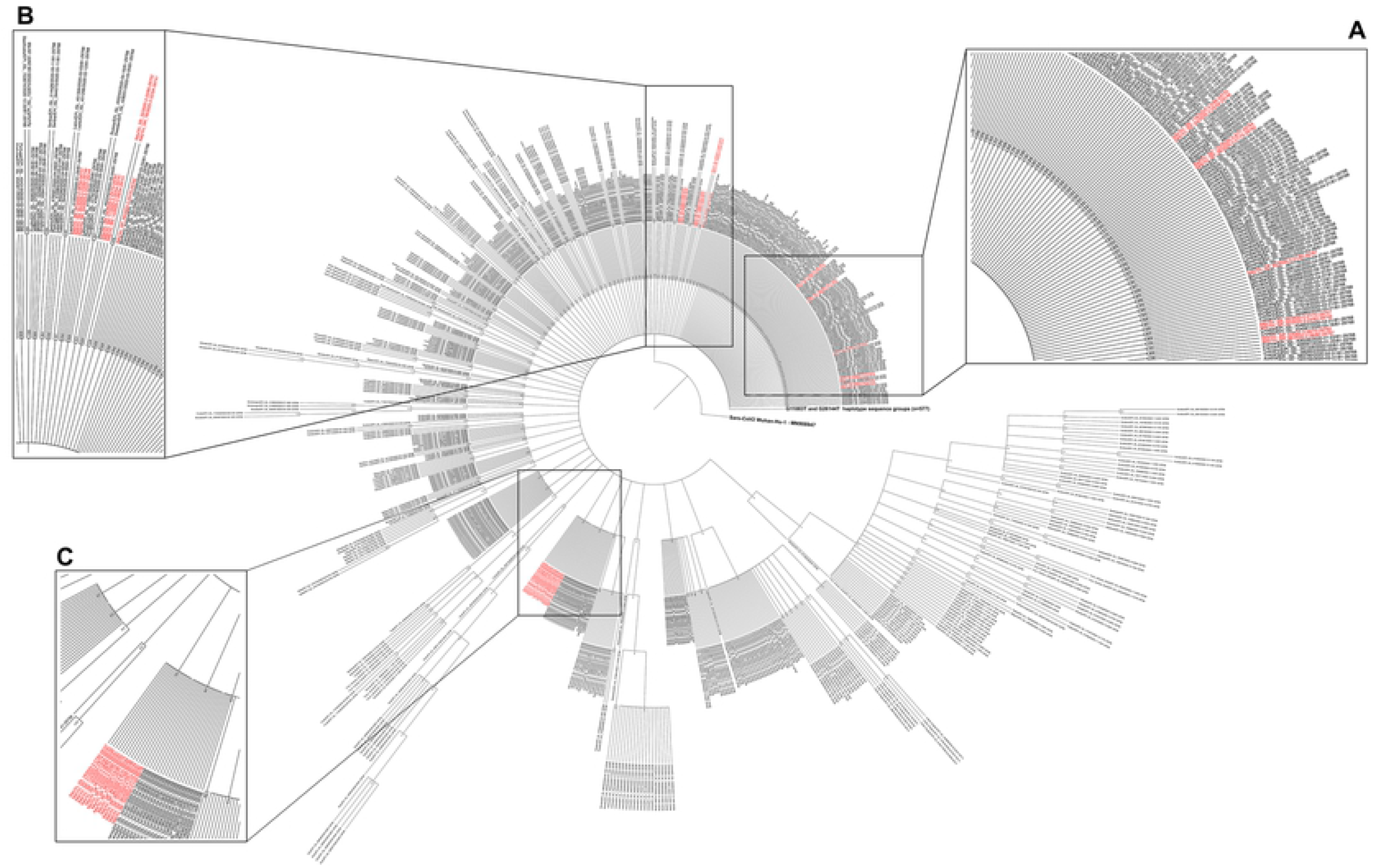
ML phylogenetic tree of European lineage B SARS-CoV-2 sequences at the beginning of the pandemic. Maximum likelihood phylogenetic tree of all European sequences collected at the beginning of the pandemic and uploaded in GIDSAID. Vo’ sequences are coloured in red. All sequences carrying the Ancestor Haplotype are collapsed. A, B and C panels zoom in the subtypes of the ancestor haplotype observed in Vo’.

**S4 Fig.**
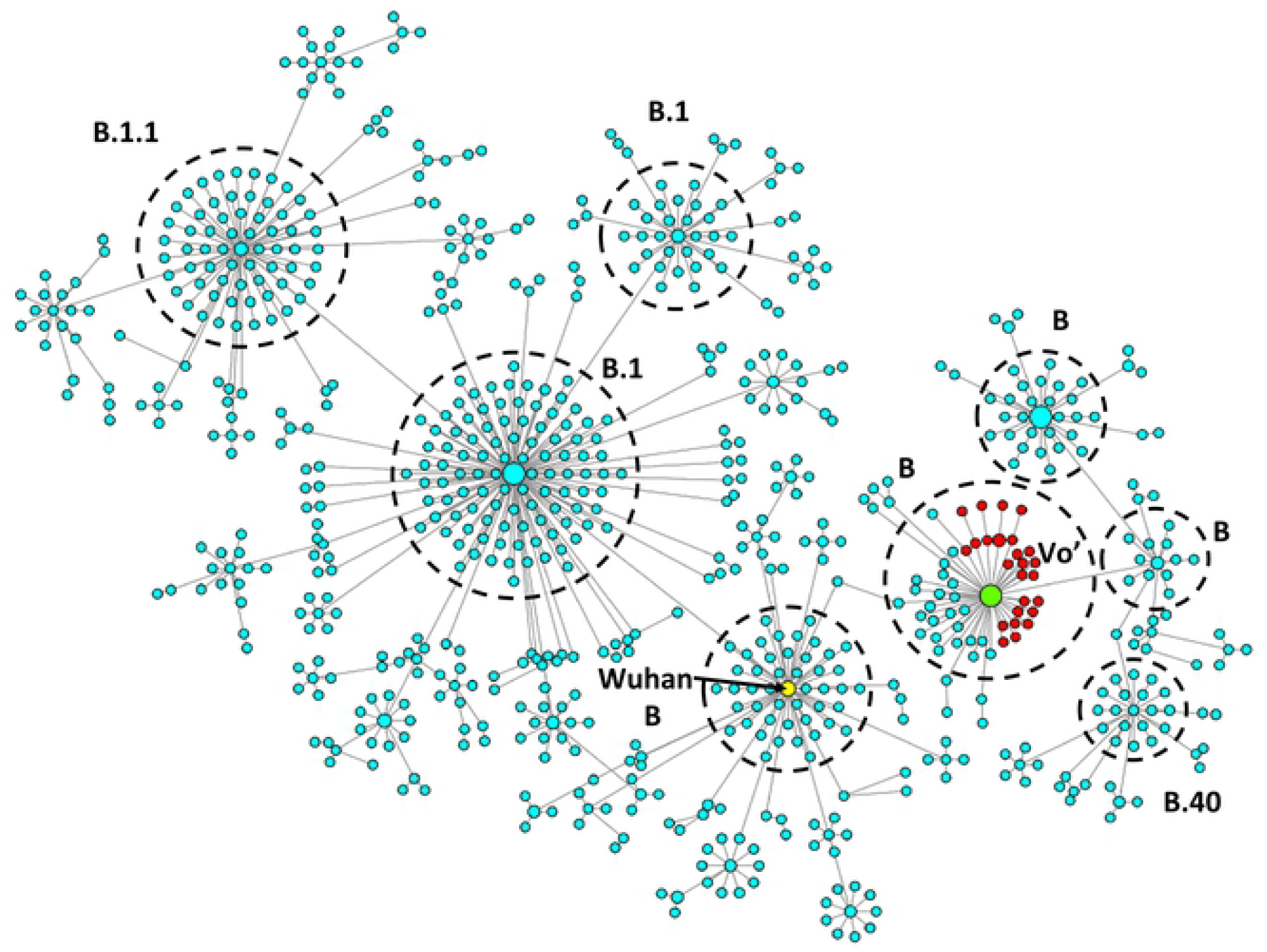
Minimum Spanning Network of European lineage B SARS-CoV-2 sequences at the beginning of the pandemic. MSN of all the European sequences collected at the beginning of the pandemic and uploaded in GISAID. Each dot represents a unique haplotype, with Vo’ haplotypes coloured in red. Haplotypes are clustered according to the lineage they belong to, with each cluster being circled with a black dotted line, and showing the basic haplotype in the central node.

**S5 Fig.**
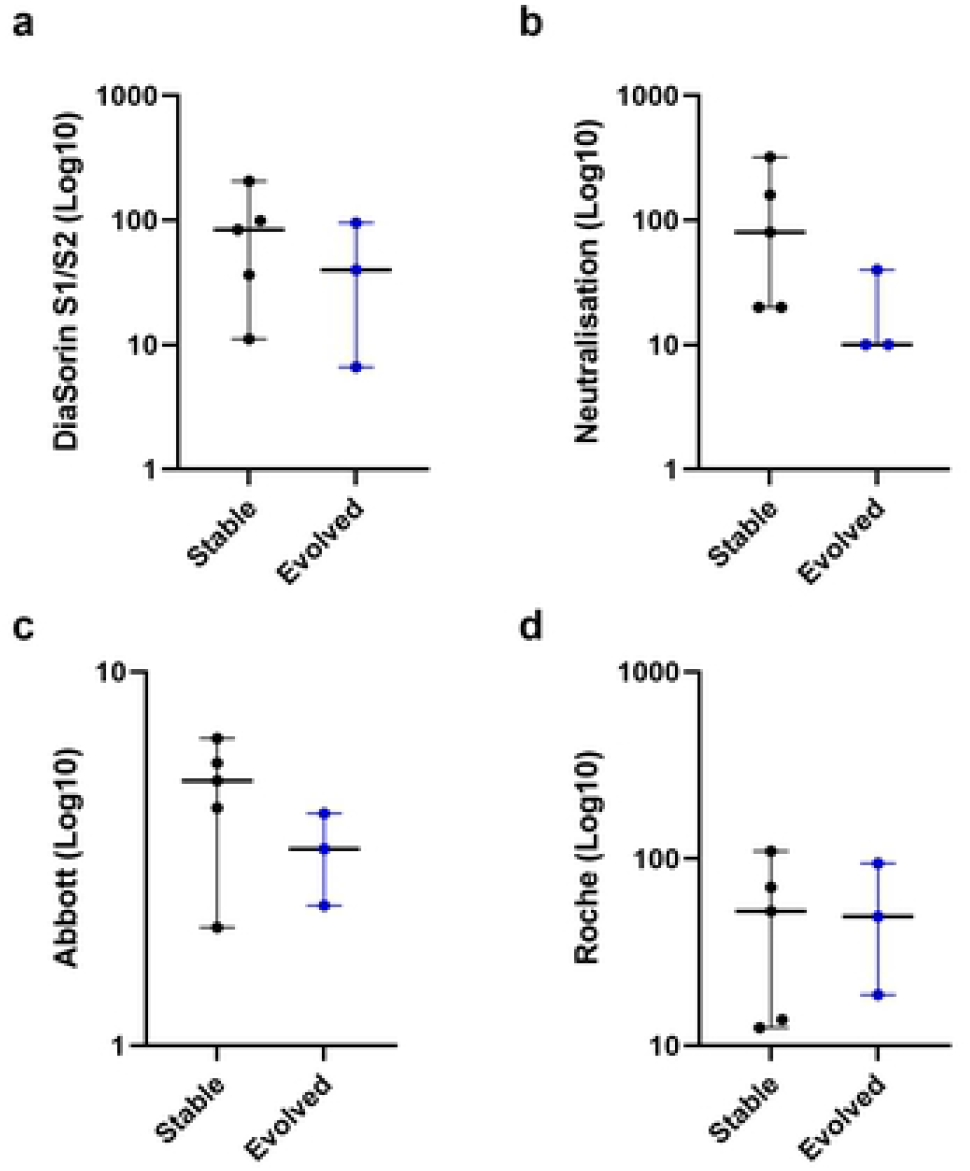
Antibody titres of subjects positive to two sequential swab tests with an average time interval of 11 days. Antibody titres observed in subjects with stable (black) and unstable (blue) viral genome in an average time span of 11 days according to DiaSorin, Neutralisation, Abbott and Roche assays (Mann Whitney test).

**S6 Fig.**
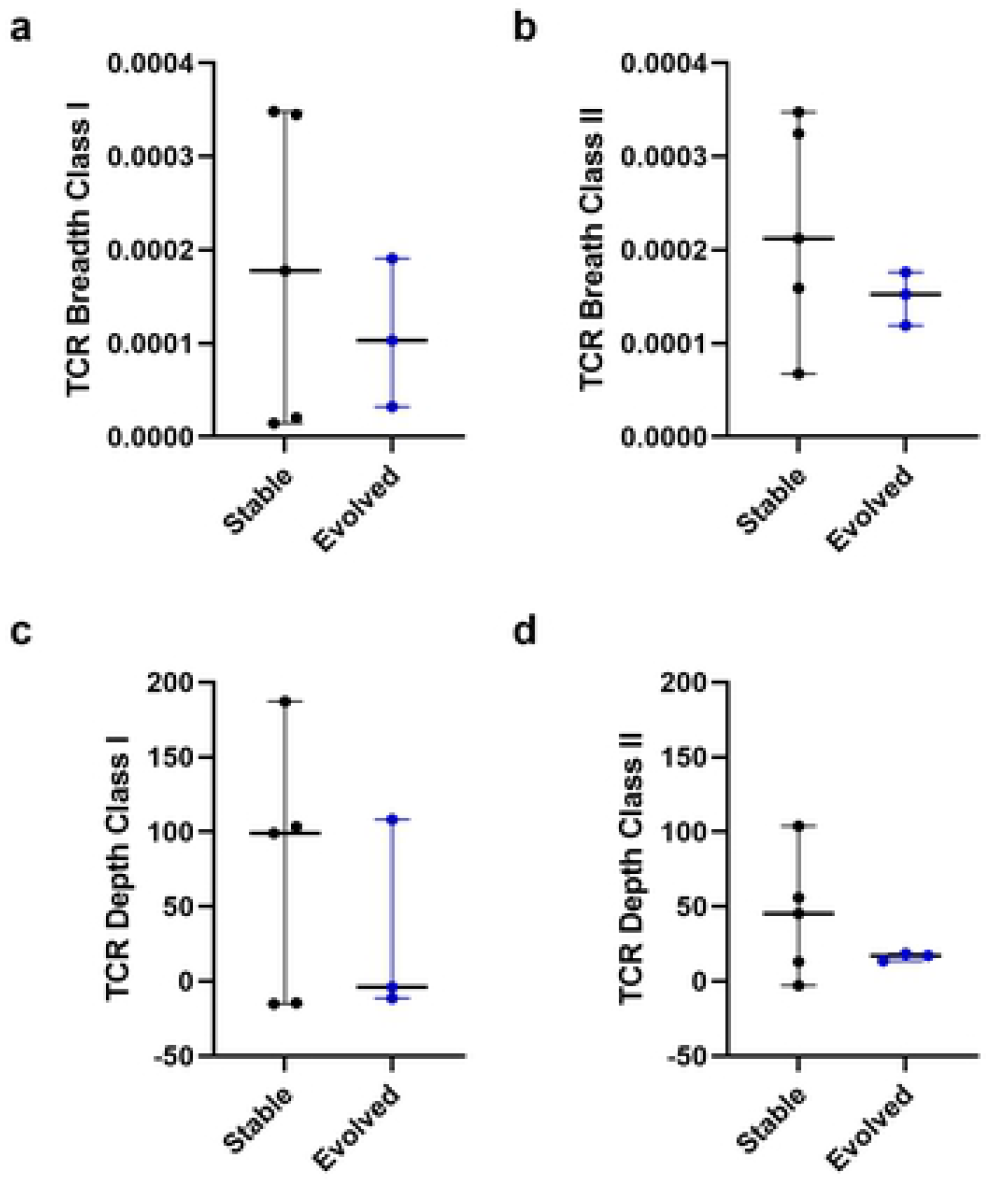
TCR features of subjects positive to two sequential swab tests with an average time interval of 11 days. TCR breadth class I (a) and class II (b) of patients with stable (black) and unstable (blue) viral genome in an average time span of 11 days (Mann Whitney test). TCR depth class I (c) and class II (d) of patients with stable (black) and unstable (blue) viral genome in an average time span of 11 days (Mann Whitney test).

**S7 Movie. Expanding Network based on symptoms onset data**

Movie based on symptoms onset data reported by the infected individuals regardless of the availability of the viral sequences. Subjects reporting contacts are depicted as rectangles, while circles represent individuals reporting no contacts with the subjects traced in the main chain. Nodes with an available sequence are coloured in green, otherwise they appear in red. Individuals who did not report a symptom onset date appear in the last frame all together.

